# Simulating the human tumor microenvironment in colorectal cancer organoids *in vitro* and *in vivo*

**DOI:** 10.1101/822684

**Authors:** Mahesh Devarasetty, Anthony Dominijanni, Samuel Herberg, Ethan Shelkey, Aleksander Skardal, Shay Soker

## Abstract

The tumor microenvironment (TME) plays a significant role in cancer growth and metastasis. Bioengineered models of the TME will advance our understanding of cancer progression and facilitate identification of novel anti-cancer therapeutics that target TME components such as extracellular matrix (ECM) and stromal cells. However, most current *in vitro* models fail to recapitulate the extensive features of the human tumor stroma, especially ECM architecture, and are not exposed to intact body physiology. On the other hand, *in vivo* animal models do not accurately capture human tumor architecture. Using the features of biopsied colorectal cancer (CRC) tissue as a guide, we address these deficiencies by creating human organoids containing a colonic stromal ECM layer and CRC spheroids. Organoids were studied *in vitro* and upon implantation in mice for 28 days. We show that the stromal ECM micro-architecture, generated *in vitro*, was maintained *in vivo* for at least 28 days. Furthermore, comparisons with biopsied CRC tumors revealed that organoids with orderly structured TMEs induce an epithelial phenotype in CRC cells, similar to low-grade tumors, compared to a mesenchymal phenotype observed in disordered TMEs, similar to high-grade tumors. Altogether, these results are the first demonstration of replicating the human tumor ECM architecture in biofabricated tumor organoids under *ex vivo* and *in vivo* conditions.

## Introduction

Tumors are products of their environment. They send signals that can have significant effects on local tissue, and they receive signals from nearby cells and extracellular matrix (ECM) that can alter their progression.^1–3^ Despite the importance of a tumor’s environment, current strategies for prognostication of tumors are centered around analyses of the tumor cells in isolation, such as morphological assessment or proliferative index calculation. Although these metrics are correlated to tumor progression, they do not capture the dynamics between a tumor and its surrounding space leading to inaccuracies when attempting to predict tumor progression and chemotherapeutic response. New technologies that improve prognostication will have a significant effect on patient mortality and lead to development of novel therapeutics which target and control tumor cells specifically, sparing healthy tissue from the deleterious effects of contemporary chemotherapeutics.

Recent studies have identified the tumor microenvironment (TME) as a major contributing factor to cancer development and growth. The combination of paracrine factors, stromal cells such as endothelial, macrophages and tumor associated fibroblasts (TAFs), ECM proteins, and tissue mechanics coalesce into the perfect environment for a cancer to thrive and evade treatment. ^2,3^ Of particular interest to our research: a number of studies have shown that tissue biomechanics^4–6^ and ECM architectures^7,8^ can alter and guide cancer cell phenotype, as well as alter therapeutic response. These findings indicate the TME as a potential target for innovative anti-cancer cell, or cancer-modulating, therapies. To devise TME targeting therapies, *in vitro* models of the TME need to be developed and validated. Many groups, including ours, have used tissue engineering techniques to fabricate tumor organoids that replicate the unique combination of factors in the TME.^9–11^ Tumor organoids can be prepared from human-sourced components yielding an accurate representation of the tumor-stroma interactions found in the TME, unlike many gold-standard animal cancer models. However, organoids cannot replicate the context of whole-body physiology to test side-effects or pharmacodynamics and pharmacokinetics.

Previously, we developed a three-dimensional (3D) organoid of the colonic submucosa complete with the unique micro-architecture found there.^9^ When we embedded tumor spheroids composed of malignant colorectal cancer cells within these organoids, we found that the tumor cells behave radically different depending on the organization of the collagen ultrastructure. In ordered, organized TMEs, CRC cells exhibited behaviors akin to healthy colonic epithelial cells with polarization and low proliferation rates. Interestingly, when we placed these same cancer cells into randomly assorted collagen I matrices, the CRC cells became highly motile and invasive with a high index of proliferation – in other words, they assumed an aggressive cancer phenotype. Furthermore, structured TMEs induced chemoresistance in CRC cells while randomly organized TMEs caused chemosensitivity.^9^ These results indicated that the presence of healthy stromal cells, capable of structuring the tumor ECM, has a suppressive effect on tumor cell phenotype and growth. Interestingly, most published data in the past decade has suggested an opposite effect of the stromal cells’ involvement on cancer progression, primarily through paracrine factors that enhance tumor cell proliferation and migration and decrease their responsiveness to chemotherapeutic drugs.^12–14^ However, these studies have typically employed TAFs, rather than normal stromal cells, and have not utilized organotypic models.^2,11,15,16^

To further our understanding of TME architecture and its role in modulating tumor growth, we analyzed CRC biopsies finding significant changes in ECM organization. Based upon our clinical observations, we fabricated CRC organoids containing hepatic stellate cells to replicate the stromal cell content and organization found in liver, the most common site of CRC metastasis.^17^ We hypothesized that the presence of stromal cells in the TME will drive ECM organization and subsequently modulate CRC tumor growth in these organoids. To broaden the application of tumor organoids as an *ex vivo* model of tumor growth, we implanted them subcutaneously in nude mice. We hypothesized that stromal cell-driven TME architecture *ex vivo* will be preserved for an extended time *in vivo*. To test our hypotheses, we analyzed TME organizations and corresponding tumor cell phenotypes in organoids for 4 weeks *in vitro* and *in vivo* and found that organoids with orderly structured stromal ECM, generated *in vitro*, maintained these structures *in vivo* throughout observation, and induced an epithelial phenotype in CRC cells. In contrast, disordered ECM allowed for mesenchymal phenotype. These results indicate that a pre-structured TME maintains its architecture in the context of whole-body physiology. Together, we present data on interactions between ECM architecture and cancer cell phenotype in three different systems, *in vitro*, in mice and in clinical samples. These findings demonstrate the clinical relevance of our CRC organoid and can be used as a model for studying tumor growth ex-vivo and prediction of potential response to chemotherapeutic drugs.

## Results

### Tumor tissue has a fewer collagen-rich areas and disorganized collagen architecture compared to normal colon tissue

The amount of collagen in a tissue is often compared between healthy and diseased tissues as a measure of disease progression. To qualitatively assess differences in the patterns of collagen-rich areas between healthy colon and CRC tissue, we performed trichrome staining. ^18–20^ We obtained colon tissue biopsies from 12 healthy, 6 low-grade, and 6 high-grade CRC patients. Healthy tissue (Fig. 1a) shows distinct compartmentalization of collagen (blue signal) outside the crypt structures of the colon, and the collagen also appears striated and aligned within the submucosal layers. CRC tissue (Fig. 1b-c), conversely, shows less collagen overall, further decreasing from low grade to high grade, and the collagen becomes more dispersed with increasing grade. Due to the differences between tissue compartments, we divided the healthy and cancerous colon tissue into two distinct areas: the mucosa/crypt and the submucosa (Fig. 1d-f). To further characterize collagen fiber organization in these healthy colon and CRC specimens, we captured the birefringent collagen signal from picrosirius red (PSR)-stained sections corresponding to those areas (Fig. 1d-f **outsets**). PSR images demonstrate a similar pattern to trichrome: aligned and bundled collagen in healthy samples, and fibrillar, disordered collagen in diseased samples.

**Figure 1:**
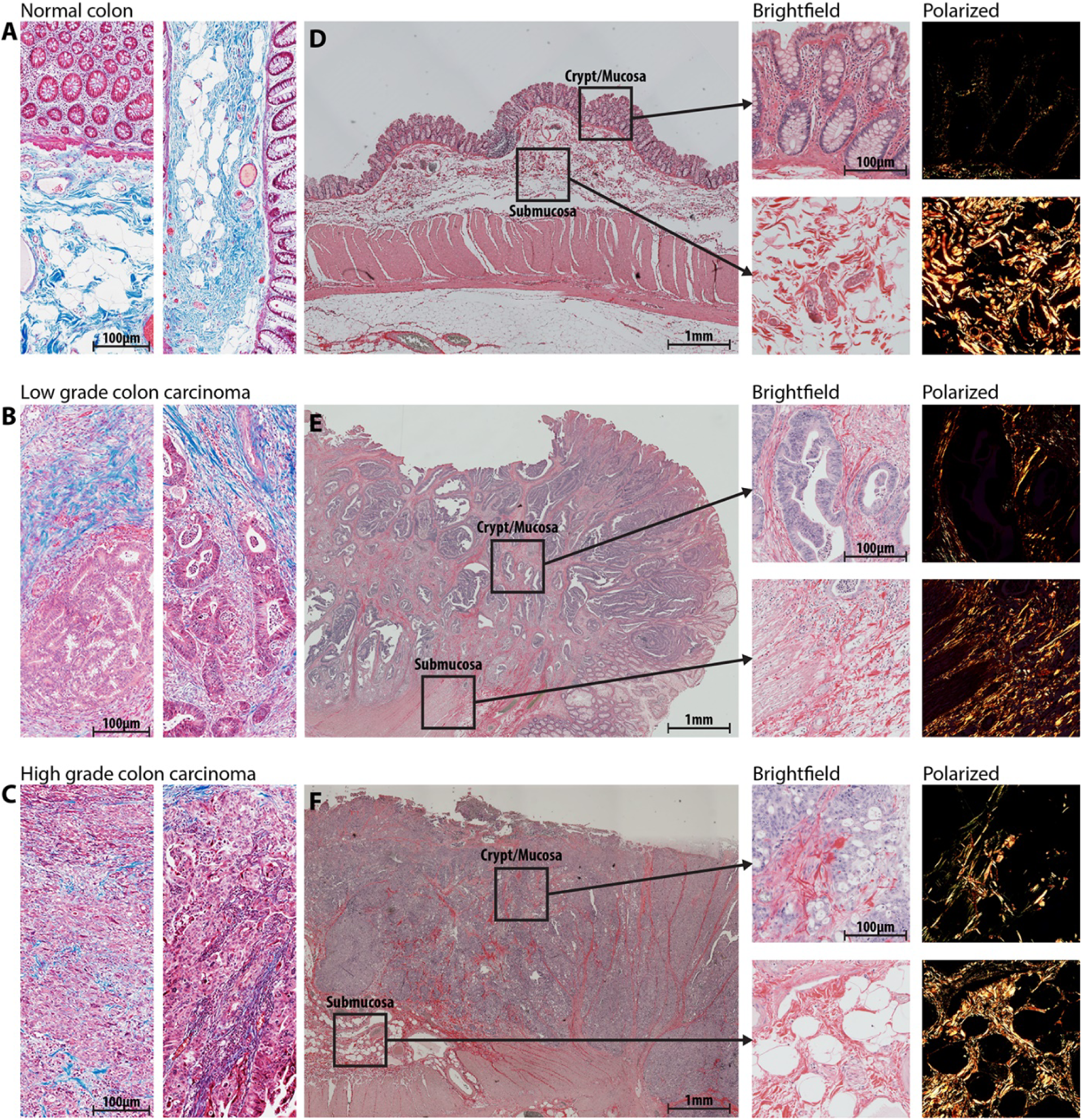
Clinical colorectal cancer sample morphology, collagen structure, and compartments. Clinical samples of varied grade, as indicated, were obtained initially stained with Masson’s Trichrome (**a,b,c**) to visualize collagen (blue signal) structure, density, and localization. H&E staining and imaging was used to qualitatively separate the tissue into mucosal and submucosal compartments (**d,e,f**). Then, PRS staining was performed and imaged to isolate collagen fibers specifically.

We digitally separated images described in Fig. 1d-f into the crypt or submucosa compartments while retaining the original tumor grade information. Then we performed automated fiber segmentation using CT-FIRE^21^ to further quantify our observations of structural changes to collagen in CRC tissue dependent on grade. In the crypt compartments, we observed that collagen fibers (Fig. 2a) become significantly more random (angular analysis) and wider in specimen derived from low- and high-grade CRC patients compared with healthy tissue (Fig. 2b-**left and middle, respectively**), whereas the fiber length decreased only in high-grade CRC patient samples (Fig. 2b-**right**). Color of PSR signal indicates the bundling or maturity of collagen: red/orange signal corresponds to bundled and thickened fibers (mature) while yellow/green labels thin and reticular collagen (young or immature). To quantify the color, we performed hue analysis with a custom Matlab script. Hue analysis of birefringent signal showed no significant differences between any of the groups (Fig. 2c). In the submucosal compartments, we found that collagen fibers had narrower angular distribution in specimen from the cancerous tissues compared with healthy tissue (Fig. 2e-**left**). Fiber width decreased in specimen from healthy individuals to low-grade and further decreased in high-grade CRC specimen (Fig. 2e-**middle**). Fiber length was lower in specimen from low- and high-grade CRC patients compared to tissue from healthy individuals (Fig. 2e-**right**). Qualitative image analysis corroborated these results showing longer and wider fibers in the healthy specimen, and a higher density of short and thin fibers in the tumor specimen (Fig. 2d). However, hue analysis of birefringent signal did not show significant differences between any of the groups, similar to the findings in the crypt (Fig. 2f). Together, these results demonstrate that collagen fiber micro-architecture changes during cancer progression with different dynamics for each colon tissue region, the mucosa or submucosa.

**Figure 2:**
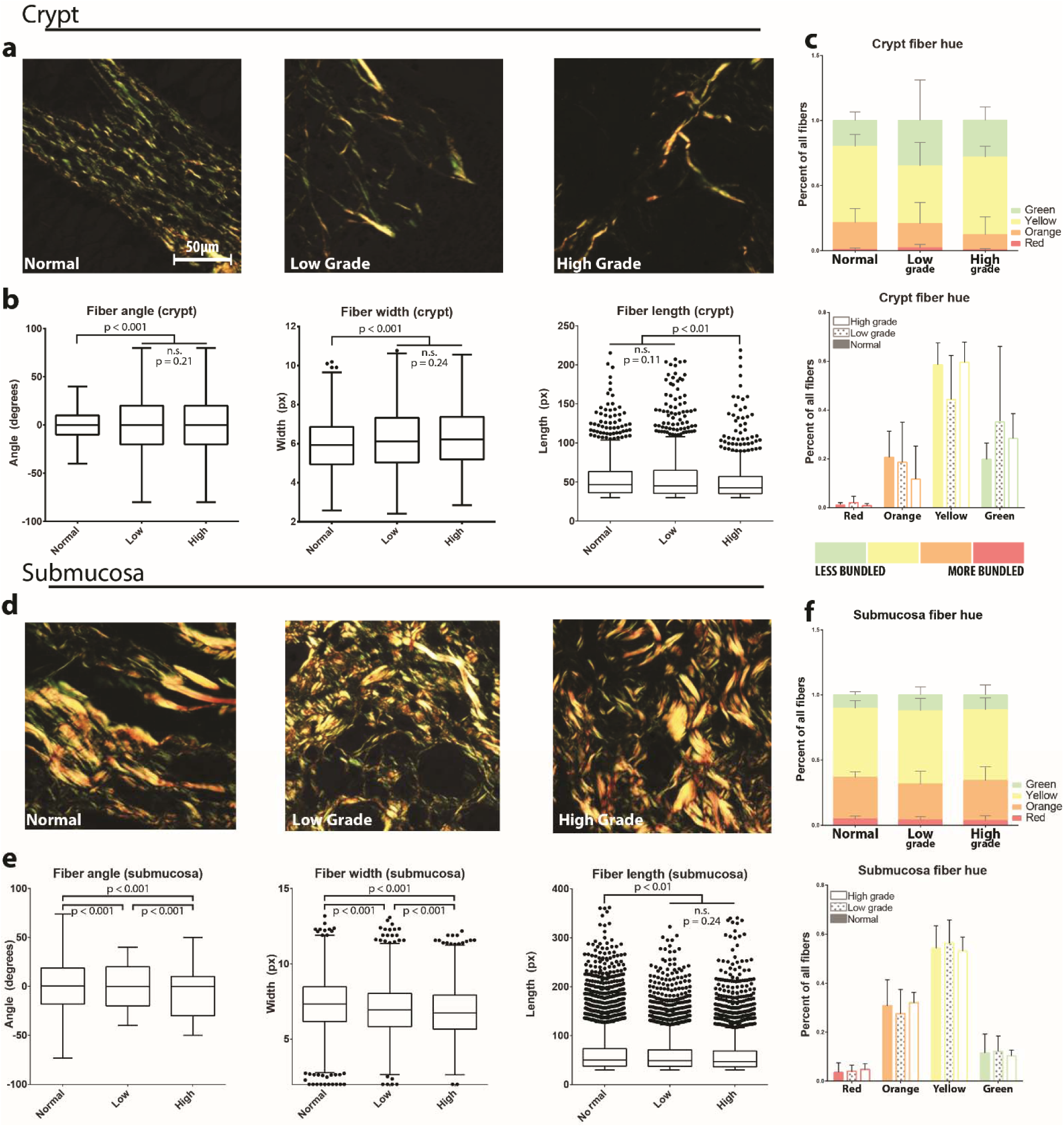
Microarchitecture of clinical samples. Regions of interest (ROIs) from the crypt and submucosal compartments of clinical samples of varying grade (n>10 ROIs from n=4 individual samples for each condition), as indicated, were PRS stained and imaged under polarized light (**a,d**). Fiber architecture was analyzed with segmentation software (CT-FIRE) to generate distributions of fiber angle, width, and length (**b,e**). Hue signal of PRS images was quantified for comparison (**c,f**). Graphs of fiber hue represent mean + s.e.m. of experiments performed in triplicate. Graphs of fiber angle, length, and width are box and whisker plots with Tukey formatting of pooled fibers from experiments performed in quadruplicate, representing 500-4000 fibers in total; individually drawn points lie beyond 1.5 * inter-quartile range of the plot.

### Replicating native collagen architectures with bioengineered organoids

To model the effects of collagen architecture on tumor phenotype, we produced tissue equivalent organoids with microenvironmental collagen architecture similar to human biopsies (outlined in **Suppl. Fig. 1a**).^9^ We embedded immortalized hepatic stellate cells (LX2) into a collagen I hydrogel which, in accordance with our previous studies,^9^ resulted in the active remodeling of collagen I and formation of structured and bundled fibers that resemble those observed in native tissue. Stellate cells participate in producing the pre-metastatic niche for CRC-to-liver metastasis and thus a represent a relevant cell type for *in vitro* CRC models. In parallel, we produced bare collagen organoids which would replicate the disorganized collagen features of high-grade CRC tumors.^9^ To test if the organoids can recapitulate the collagen organization features observed in healthy colon, low-, or high-grade CRC, we characterized collagen fiber bundling and micro-architecture in organoids cultured *in vitro* and in organoids implanted in mice. An additional goal was to determine if organoids formed *in vitro* could retain collagen fiber organization under physiological conditions *in vivo*. Similar to the analysis of collagen fiber organization in the clinical specimens (Fig 2), we used PSR staining and image segmentation tools to assess collagen fiber features in the *in vitro* cultured and *in vivo* explanted organoids (Fig 3).^21^ Visual inspection of the stained collagen fibers in the organoids showed that the LX2 organoids produce increased red and orange signal *in vitro* and *in vivo* compared with the collagen-only organoids (Fig. 3a). Hue analysis corroborated these findings and showed that LX2 organoids cultured for 7 days *in vitro* produced bundled collagen fibers with higher levels of orange and yellow collagen (Fig. 3b) suggesting that these samples were more organized compared with collagen-only organoids. Similarly, organoids explanted after 14 days showed that LX2 organoids possessed higher degrees of bundling with significantly higher amounts of red and orange signal compared with collagen-only organoids.

**Figure 3:**
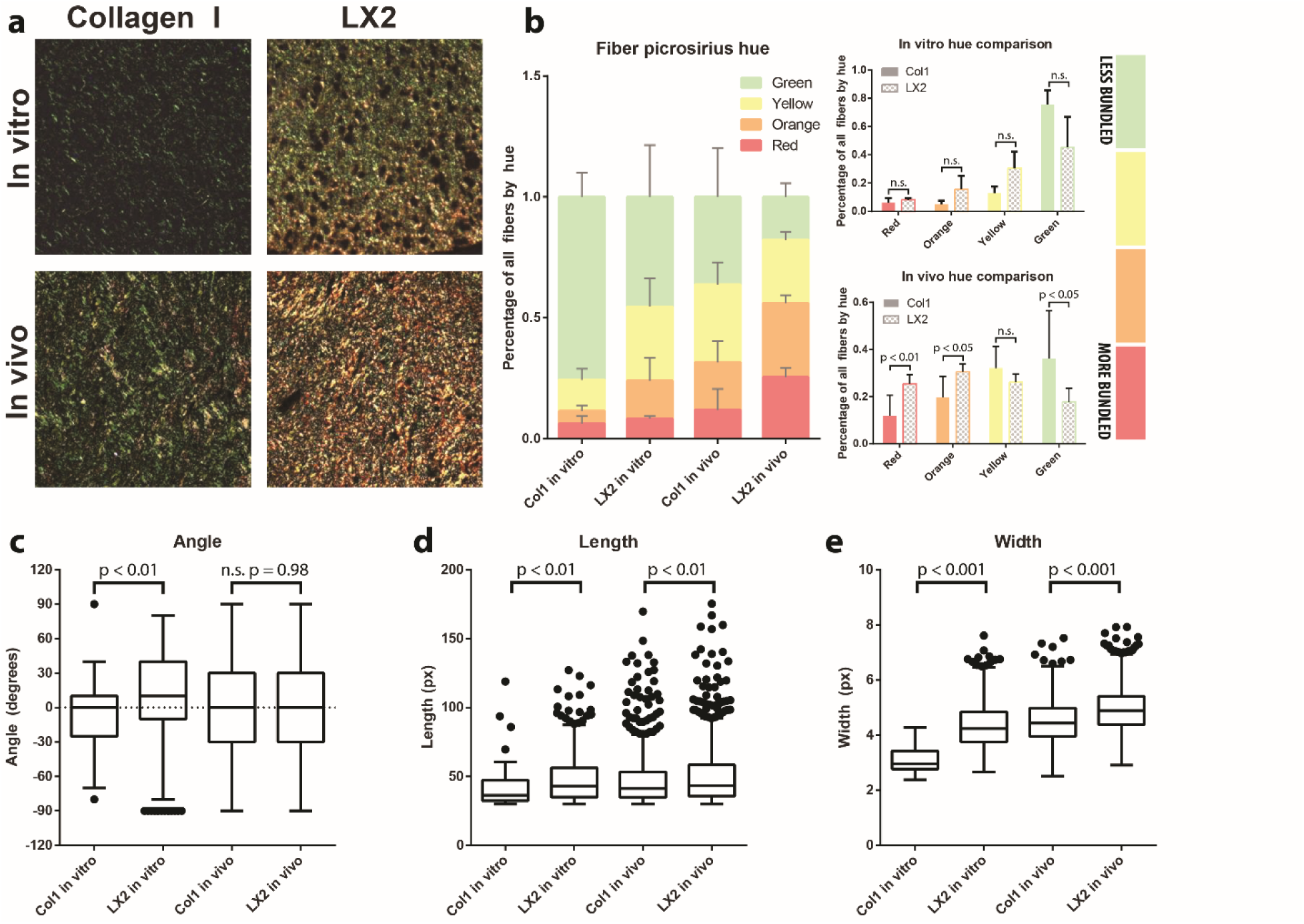
Organoid extracellular matrix (ECM) organization *in vitro* and *in vivo*. Collagen fiber microarchitecture in collagen control and LX2 organoids, *in vitro* and *in vivo*, as indicated, was visualized with PRS (**a**). Fiber bundling was quantified through signal hue analysis (**b**): green and yellow signal indicate less bundled fibers and orange and red signal indicate more bundling. Collagen fibers were quantified with segmentation software (CT-FIRE). Distributions of angle (**c**), length (**d**), and width (**e**) of fibers were obtained from organoids *in vitro* and *in vivo*. Graphs of fiber hue represent mean + s.e.m. of experiments performed in triplicate. Graphs of fiber angle, length, and width are box and whisker plots with Tukey formatting of pooled fibers from experiments performed in quadruplicate or greater; individually drawn points lie beyond 1.5 * inter-quartile range of the plot.

Quantitative analysis of collagen fiber organization showed a wider angular distribution in LX2 organoids compared with collagen-only organoids (Fig. 3c). However, in *in vivo* explanted organoids no significant difference was observed in the angular distributions of collagen fibers between groups, suggesting a similar, random, assortment of fibers overall (Fig. 3c). On the other hand, fiber length and width were significantly higher in the LX2 organoids *in vitro* compared with collagen-only organoids and this trend was similar for *in vivo* explanted samples (Fig. 3d & e). Together, these results suggest that LX2 remodeling of collagen *in vitro* generates more bundled and ordered fiber micro-structures and importantly, these ordered collagen fiber features were maintained after 14 days of *in vivo* implantation. Furthermore, the differences in collagen fiber organization between LX2-containing organoids and collagen-only organoids simulate differences between healthy colon and low-, or high-grade CRC, respectively.

### Embedded tumor spheroids grow slower in organized microenvironments

Next, we tested the effects of the microenviromental (collagen fiber) organization on tumor cells present within organoids with distinct collagen fiber features. We inserted spheroids of HCT116, metastatic CRC cells, into both LX2-contaning and collagen-only organoids (**Suppl. Fig. 1b**) and analyzed tumor cell phenotype in *in vitro* cultured and *in vivo* implanted organoids.

H&E-stained images show that HCT116 spheroids inside the LX2 organoids remained compact and were significantly smaller than the collagen-only organoids after 7 days (Fig. 4ai-lower, c), and continued to decrease in size and retained a compact, epithelial-like morphology by 28 days. In contrast, HCT116 cells embedded in collagen-only organoids migrated outside the initial spheroid body, and after 7 days, protrusions of cells were apparent in most radial directions. This migratory behavior continued and resulted in a significant increase in aggregate size after 28 days (Fig. 4ai-upper, c), and loss of compact morphology, possibly due to a mesenchymal phenotype induced by the unstructured matrix organization, as we have previously shown.^9^ Progressive measurements of HCT116 spheroid growth over time indicated that spheroids in LX2 organoids were size restricted, and even exhibited size reduction, whereas HCT116 spheroids in collagen-only organoids were able to expand (Fig. 4c).

**Figure 4:**
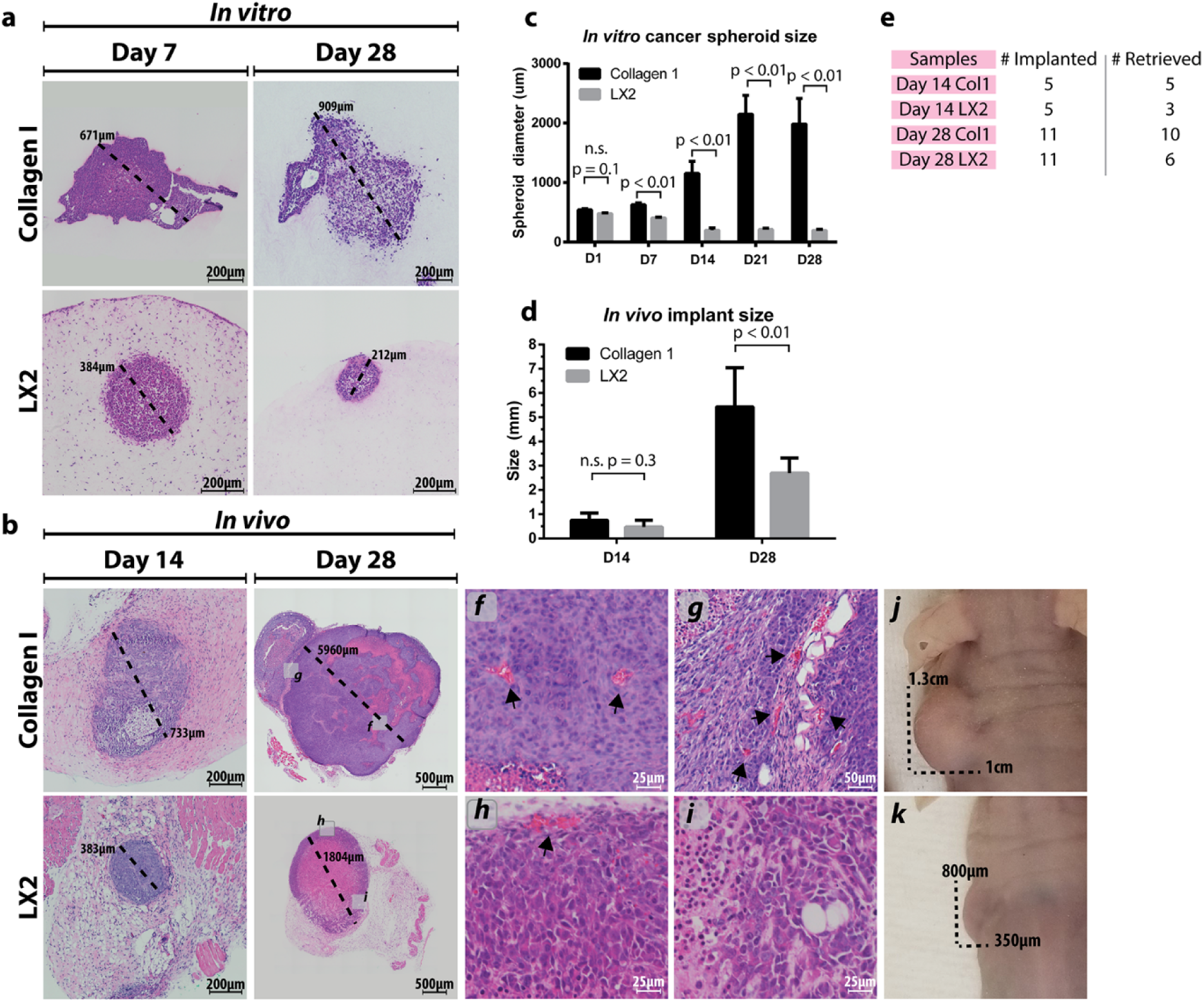
Tumor organoid morphology and growth *in vitro* and *in vivo*. Tumor spheroid morphology was visualized on days 7 and 28 *in vitro* (**a**) and *in vivo* (**b**) in both collagen control and LX2 organoids, as indicated. Integration of vasculature was also observed at day 28 of *in vivo* implanted organoids (insets **f,g,h,i**). Spheroid diameter was tracked over time *in vitro* (**c**), and *in vivo* implanted organoids were measured at the time of explantation for average diameter (**d**). Examples of gross implant size *in vivo* can be seen in insets **j** & **k**. Implant retrieval data was also collected for *in vivo* studies and collected (**e**). Graphs represent mean + s.e.m. of experiments performed in triplicate.

Next, we analyzed the size and morphology of *in vivo* implanted LX2 and collagen-only organoids after 14 and 28 days (Fig. 4b). H&E-stained images indicate that HCT116 spheroid growth in LX2 organoids was restricted at day 14, and a large necrotic core had formed by day 28, although the tumor compartment had continued to grow in size. Blood vessels were evident at the perimeters of the tumor compartment in the LX2-containing organoids, suggesting that the tumor cells failed to induce neovascularization inside the tumor mass (Fig. 4b **inset** h & i). Conversely, HCT116 spheroids in collagen-only organoids grew rapidly and by day 28, the majority of the HCT116 tumor compartment appeared viable with a low proportion of necrotic area and extensive neovascularization in the tumor mass (Fig. 4b-**upper and inset** f & g). Overall, collagen-only explanted organoids contained an average of 35 vessels compared to approximately 4 in explanted LX2 organoids (**Suppl. Fig. 2**). Size measurements of explanted organoids showed no differences in average implant diameter explanted at day 14, however LX2 organoids were significantly smaller compared to collagen-only organoids by day 28 (Fig. 4d) and significant size differences could be observed before organoid excision (Fig. 4b **inset** j & k). Of note the number of organoids that could be explanted were different between the two groups. Fewer LX2 organoids were successfully retrieved compared with collagen-only organoids at both day 14 (LX2: 60%; Col: 100%) and day 28 (LX2: 55%; Col: 91%) (Fig. 4e), suggesting that the presence of LX2 cells in the organoids represses the growth of the HCT116 spheroids and could lead to the loss of the tumor organoid tissue.

### Relationships between collagen fiber organization and architecture and cancer cell phenotype in CRC organoids and CRC tissue biopsies

Tumor spheroid growth in organoids of different collagen architectures suggested that it may simulate differences between low and high-grade cancers. We previously observed that collagen fiber micro-architecture and topography had a significant impact on EMT in CRC cells in these organoids.^9^ Accordingly, to assess phenotypic similarities between organoids and biopsied tissue, we performed immunohistochemistry (IHC) analysis on tissue sections from CRC organoids and CRC tissue biopsies for markers associated with EMT and oncogenesis (**Suppl. Figs. 3-7**) and used custom segmentation algorithms in Visiopharm^TM^ to quantify their expression.

Tumor cell proliferation was determined by staining for Ki-67 and analyzing its nuclear expression (**Suppl. Fig. 3**). In both *in vitro* cultured and *in vivo* explanted organoids, Ki-67 expression was higher in collagen-only organoids compared with LX2 organoids. Similarly, increasing Ki-67 expression was associated with higher grading in tumor samples (Fig. 5a). E-Cadherin is a cell-cell adhesion protein, present in healthy colonic epithelial cells and can be lost when cancer cells undergo EMT. E-Cadherin was found around the cell membrane and appeared continuous around positive cells in IHC images (**Suppl. Fig. 4**). We quantified E-Cadherin expression by counting cells with completely stained membranes only (Fig. 5b). *In vitro*, LX2 organoids showed significantly higher expression of E-Cadherin compared with collagen-only organoids. *In vivo*, no significant differences in E-Cadherin expression were observed between groups. Notably, CRC tumor biopsies demonstrated a significant decrease in E-Cadherin expression, compared with healthy colonic tissue and the expression further decreased with increasing tumor grade (Fig. 5b). N-Cadherin is another cell-cell adhesion protein that is typically found in colorectal cancer cells that have undergone EMT.^22^ We quantified N-Cadherin in the same manner as E-Cadherin, but also counted cells with nuclear expression of N-Cadherin as that is indicative of cytoplasmic expression that has not fully localized to the membrane (**Suppl. Fig. 5**).^22^ Collagen-only organoids displayed significantly higher levels of N-Cadherin expression compared to LX2 organoids both *in vitro* and *in vivo* (**Suppl. Fig. 4** and Fig. 5c). CRC tumor biopsies and healthy colonic tissue showed very low N-Cadherin expression overall with about 1-3% of cancer cells positive for N-Cadherin staining. The WNT/β-Catenin pathway is an important pathway associated with EMT, in which β-Catenin protein ultimately accumulates in the nucleus where it acts as part of a transcription factor complex and facilitates the upregulation of a host of oncogenic processes.^23^ We quantified β-Catenin expression by counting cells with positive nuclear-localized β-Catenin signal (**Suppl. Fig. 6**). Collagen-only organoids had higher levels of nuclear β-Catenin compared with LX2 organoids in both *in vitro* and *in vivo* groups, although the difference was not statistically significant (*in vitro:* p = 0.058) (Fig. 5d). Tumor tissue specimen derived from patients with high-grade CRC exhibited the highest levels of nuclear β-Catenin expression, corroborating the previous results that higher grades of CRC are associated with EMT. The focal adhesion kinase (FAK) pathway is also associated with oncogenesis and EMT. FAK is a tyrosine-kinase that forms adhesions with matrix components, and high levels of FAK are found in aggressive cancer cells, whereas knockdown of FAK can reduce or eliminate cancer cell motility.^24^ We quantified FAK by measuring the positively stained area fraction of the total tumor area (**Suppl. Fig. 7**). FAK expression was significantly higher in collagen-only samples compared with LX2 organoids in both *in vitro* and *in vivo* groups (Fig. 5e). Interestingly, low-grade tumor specimen demonstrated the highest FAK expression while high-grade tumor specimen had increased expression compared to normal colonic tissue, but a lower expression compared to low-grade. This result suggests that in collagen-only organoids cancer cell motility could be enhanced, similar to human tumors. Together, expression analysis of this selected protein panel indicates that tumor cell spheroids embedded in collagen-only organoids showed an average phenotype that correlates with a more advanced grade of CRC, compared with spheroids embedded in LX2 organoids which appear to have a phenotype correlated with low grade CRC.

**Figure 5:**
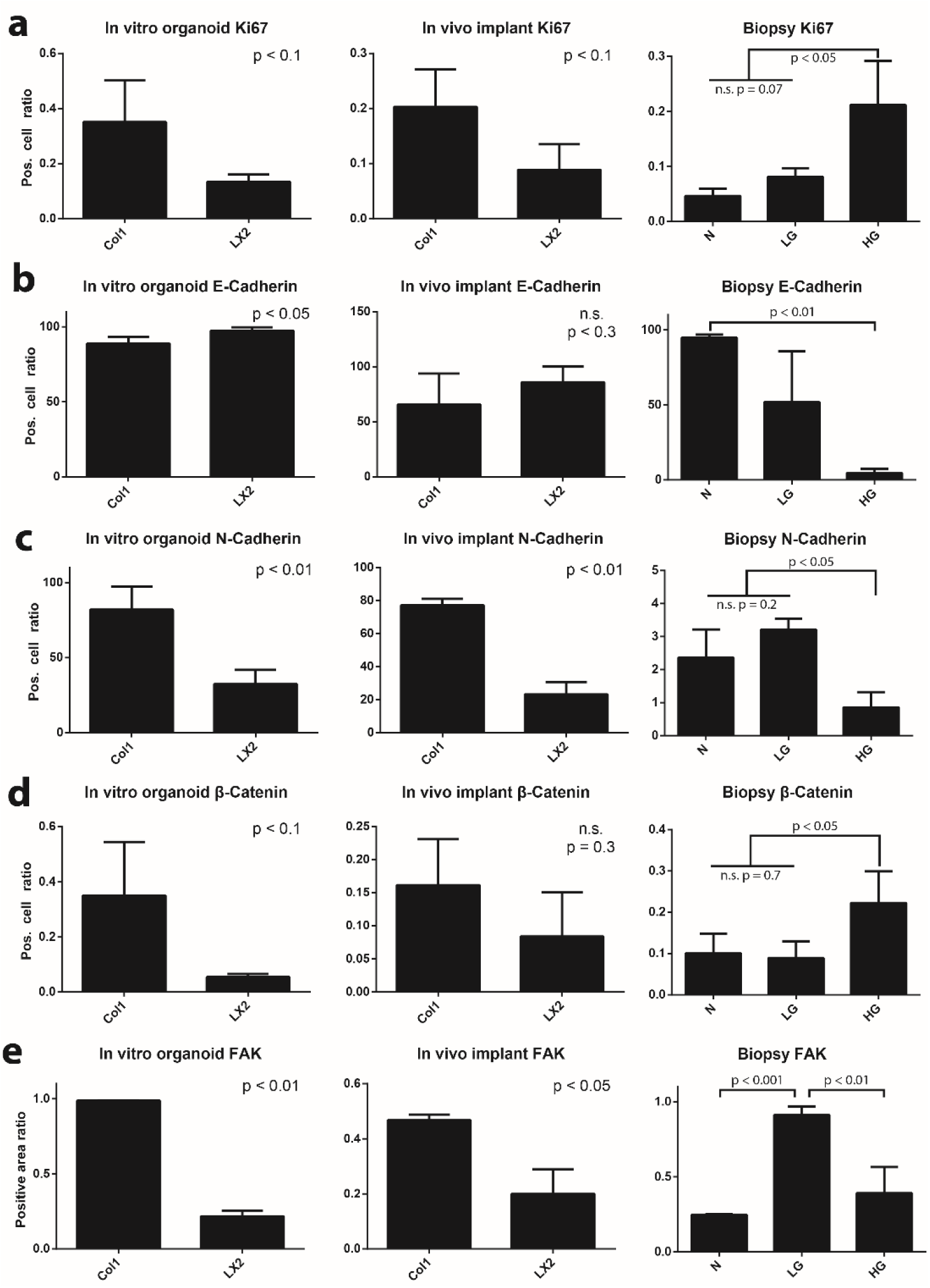
Immunophenotyping of tumor cells within organoid and clinical sample. Samples from organoids cultured *in vitro*, organoids implanted *in vivo*, and clinical CRC biopsies of varying grade were immune-stained for markers related to EMT and oncogenesis, as indicated. Staining results were analyzed using Visiopharm and graphed as the proportion of cancer cells with: (**a**) nuclear localization of Ki67; (**b**) with fully intact, membrane localized E-Cadherin expression; (**c**) with N-Cadherin expression; (**d**) with nuclear localization of β-Catenin. (**e**) Ratio of total area corresponding to positive FAK expression in tumor spheroid and clinical CRC biopsies of varying grade. Graphs represent mean + s.e.m. of three regions of interest from each sample of experiments performed in triplicate.

### Tumor spheroids grown in unorganized microenvironments are more sensitive to chemotherapeutics

Since the HCT116 tumor spheroids demonstrated significantly different phenotypes in LX2 and collagen-only organoids, we speculated that they will also have different responses to clinically used chemotherapeutic agents. To test this hypothesis, we exposed tumor spheroid bearing organoids to 4 drugs: 5-Fluorouracil (5-FU), which disrupts thymidylate synthase, and its clinically-used drug combinations FOLFOX and FOLFIRI, and a targeted therapy, Regorafenib. FOLFOX is a 5-FU combination with folic acid (leucovorin) and oxaliplatin (platinum-based antineoplastic). FOLFIRI is a combination of 5-FU, folic acid, and irinotecan (topoisomerase inhibitor). Regorafenib is a multi-receptor tyrosine kinase inhibitor.

Organoids were treated for 72 hours, then harvested and stained for Ki67 and Caspase 3 in order to determine the proportion of cells that were either actively proliferating, apoptotic, or neither (Fig. 6a & b). All chemotherapeutics had minimal effect on apoptosis of tumor cells within an LX2-organized stroma. In contrast, HCT116 spheroids in collagen-only organoids were highly sensitive to all 5-FU and its combinations and displayed significantly higher levels of apoptotic cells compared to LX2 organoids. On the other hand, all chemotherapeutics arrested the proliferation of the tumor cells in both LX2 and collagen-only organoids. However, the effects of 5-FU and FOLFOX on tumor cell growth arrest was significantly higher in collagen-only organoids compared with LX2 organoids. Interestingly, the effects of FOLFIRI treatment on growth arrest was significantly higher in LX2 organoids compared with collagen-only organoids. Finally, Regorafenib induced a significant growth arrest in tumor cells, but with little difference between LX2 and collagen-only organoids.

**Figure 6:**
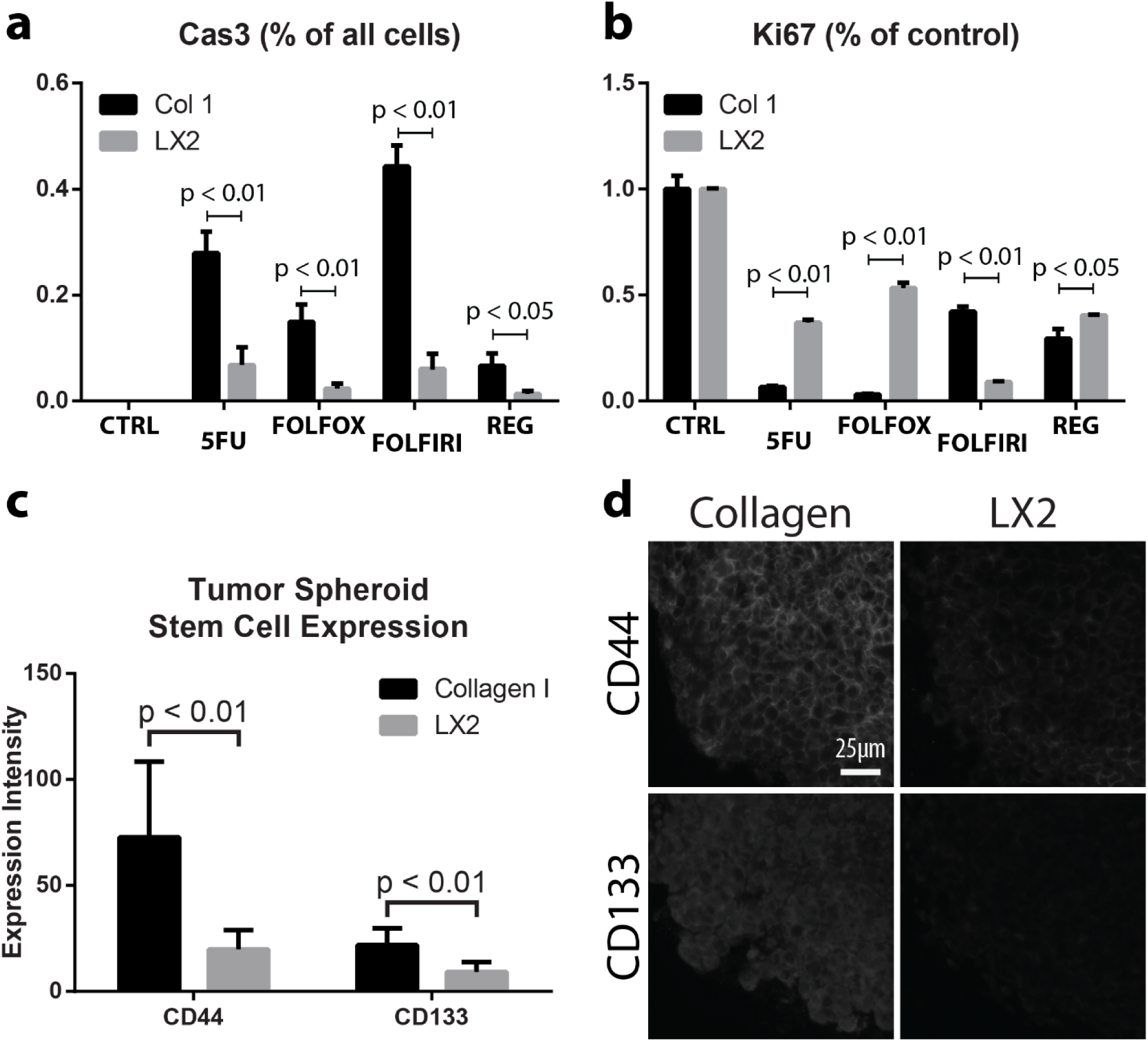
Chemotherapeutic sensitivity and expression of cancer stem cell markers in CRC organoids. Collagen control and LX2 organoids, as indicated, were exposed to various chemotherapeutics for 72 hours, and the expression of Caspase3 (measured as percent of all cells) (**a**) and Ki67 (measured as percent of control) was analyzed (**b**). CD44 and CD133 expression was quantified in collagen control and LX2 organoids, as indicated, (**c**) using IHC (**d**). Graphs represent mean + s.e.m. of three regions of interest from each sample of experiments performed in triplicate or greater.

Cancer stem cells are often considered as less chemotherapy-responsive compared with the rest of the tumor cell population. HCT116 and other CRC cells were reported to expressed several known CSC markers, especially CD44 and CD133.^25^ To identify potential differences in stem cell markers between organoids with ordered (LX2) and disordered (collagen-only) stromas, we measured CD44 and CD133 expression in HCT16 spheroids in these organoids (Fig. 6d). CD44 was expressed in both LX2 and collagen-only organoids with significantly higher levels found in collagen-only vs. LX2 organoids (Fig. 6c). CD133 expression was lower, by comparison to CD44 expression, but similarly, was higher in collagen-only organoids compared to LX2 organoids. These results suggest that disordered, collagen-based stroma, enriches the cancer stem cell population in HCT116 spheroids embedded in collagen-only organoids.

## Discussion

In patients, tumors grow and progress over many years, often-times remaining in a non-invasive equilibrium for extended periods without producing any outward symptoms. Events, not fully understood, shift the tumor from a non-invasive phenotype towards an invasive one that is associated with rapid changes to the TME, including ECM degradation and stromal cell activation, and tumor cell migration and proliferation. Although 2D cell culture has led to many breakthroughs in cancer research, it cannot replicate the nuances of the TME which have been found to significantly affect tumor progression. Not only are tumors composed of a myriad of cell types that secrete a unique combination of signaling factors, but cancerous cells are attached to, and often-times require, the latticework of proteins that comprise the ECM – the physical TME. This means that therapeutics developed in 2D culture systems may not translate well to treating tumors *in vivo*, impacting patients as well as leading to high drug development costs. Tumor organoid technology, developed in recent years, addresses the deficiencies of traditional cell culture systems by replicating some of the unique features of the TME.^9–11^ We have recently published several studies describing how tumor organoid technology, which allows to systematically manipulate structural features of the TME and subsequently test the effects on tumor cell phenotype.^9–11^ Despite the value of 3D tumor organoids, their maintenance under artificial culture conditions *in vitro* cannot fully assess tumor cell growth *in vivo*. However, *in vivo* mouse models are poor mimics of human physiology and tumor growth kinetics, as the TME will be largely composed of murine components. The goal of this study was to transition the replicative features of *in vitro* bioengineered organoids into an *in vivo* model to capture the best of both methods. We replicated native TME architecture, as observed in clinical biopsies, with CRC organoids, then implanted them into nude mice to expose them to *in vivo* physiology. Our results demonstrate that organoids, fabricated to recapitulate the TME and phenotype of different tumor grades, maintained ECM architectural features for up to 4 weeks *in vivo*, and furthermore, CRC cells retained their original, intended grade-related phenotypes.

In the current study we used hepatic stellate cells (LX2) to simulate stromal cells that are involved in collagen remodeling of the liver metastasis microenvironment.^26–29^ The results, demonstrating that LX2-mediated ECM structured microarchitecture push the HCT116 metastatic CRC cells towards an epithelial phenotype vs. a mesenchymal phenotype induced by unstructured ECM architecture are especially intriguing as they suggest that under certain conditions the hepatic stellate cells, commonly known to support EMT and growth of liver metastasis^9,30,31^, may act to suppress growth and spreading of metastasis if given an opportunity to create structured matrix around the metastatic foci. Furthermore, HCT116 cells demonstrated reduced chemotherapeutic sensitivity in the LX2 organoids as well. Others have also targeted hepatic stellate cells (HSCs) as possible TAFs and as contributors to fostering a premetastatic environment for CRC cells. For example, addition of conditioned media from activated-HSCs to CRC cells in 2D culture resulted in increased motility and invasive characteristics of the tumor cells.^31,32^ This data suggests a paracrine effect of HSCs on the tumor cells however, these studies were performed in a non-physiologic 2D culture system, which does not capture important features of the TME, like our organoid model can. Although paracrine secretion of factors from TAFs can clearly drive cancer progression, TAF’s involvement does not stop at the level of external cell signaling; TAFs have the ability to change ECM structure and organization, and understanding the subsequent effects on cancer cells is an important step towards more accurately modeling the TAF-cancer cell axis.

A major motivation of this research is to create a better *in vitro* model of tumor tissue by including, alongside the tumor cells, stromal cells and ECM. To achieve this aim, we utilized a bottom-up approach to replicate the distinct compartments of tumor tissue, as observed in Fig. 1. The native colonic submucosa is collagen-rich and populated by a mixture of SMCs, fibroblasts, adipose and immune cells which informed our decision to utilize collagen as the base hydrogel with fibroblasts as the main stromal component. In addition, tumor compartments in biopsies were typified by aggregates (foci) of cancer cells which we simulated with a cancer spheroid. This approach replicated the dense cellularity, diffusion kinetics, and cell-cell contacts evident in human tumors. Our system is inherently modular as well; each compartment can be modified independently to replicate the tumor-stroma dynamics of a wide variety of tumor etiologies. The HCT-116 cells used here can be replaced by cells of lower malignancy, patient cells, or a heterogenous mixture of cancer cells. The stromal compartment can be modified to include a wider range of cells, higher concentrations of collagen, or different ECM components such as laminin, fibronectin, or elastin. In our previous work, we utilized colonic smooth muscle cells within the stromal compartment to replicate the native, healthy colon.^9^ We also previously demonstrated that stromal cells in our platform are responsive to biochemical manipulation, in this case to aminopropionitrile (BAPN), an inhibitor of lysyl oxidase, which decreased smooth muscle cell collagen remodeling capability. In human tumors, a panoply of paracrine signals (TGFβ1, PDGF, IL-1β, etc.) drive stromal cell involvement, many of which can be modeled within our system. In fact, exposure of tumor organoids to TGFβ1 further stimulated ECM restructuring, creating highly bundled collagen fibers and resulting in stiffer organoids (manuscript in preparation).

An important observation of this study is the consistency of results between the *in vitro* experiments and under physiological conditions *in vivo*, both of which agree with our prior findings.^9,11^ These results indicate the specific architecture of the niche may help determine EMT or MET phenotype of liver metastatic tumor cells, and could explain why some tumors do not respond well to non-combinatory, anti-proliferative chemotherapies.^33^ This highlights an important feature: the ability for our tumor organoids to demonstrate similar phenotypes in embedded tumor cells as seen in clinical tumor biopsies. Further, we can challenge our organoids with a large library of small molecule inhibitors, chemotherapeutics, and experimental compounds and expect correlation between our results and those found clinically. Future directions may include using mathematical modeling to transfer results found *in vitro* to predictive information for patient prognosis or drug response. Lastly, we have demonstrated that fiber organization and fibroblast-mediated ECM remodeling appears to affect cancer cells and is correlated with cancer grades in clinical specimens, indicating these metrics have significant potential to be used as prognostic and/or diagnostic tools.

Overall, this study is the first to characterize an *in vitro* model of the TME based on observations of native tissue and validate it in the context of whole-body physiology. Non-traditional treatment vectors that target the ECM or stromal cells might provide valuable avenues for developing novel treatments or co-therapies that synergize with existing chemotherapeutic or radiation technologies. By controlling cancer cell responsiveness through changes to the TME, lower doses of chemotherapy or radiation could become effective thereby reducing or eliminating many of the undesirable side-effects of traditional cancer therapies (*e.g.*, cytotoxicity in healthy tissues), as well as yielding lower tumor cell resistance rates. Organoids are a promising modality for drug development and screening because they can reproduce human physiology at a high level. However, understanding of pharmacokinetics and pharmacodynamics still necessitates the use of animal models that are, however, not without their own limitations when being translated to humans. Here, we bridge the gap between *in vitro* study and animal modeling by utilizing implanted human tumor organoids, and outline a novel mechanism of cancer cell control – ECM microarchitecture.

## Materials and Methods

### Cell Culture

Human hepatic stellate cells, LX2 cells, were provided by Dr. Scott Friedman (Icahn School of Medicine at Mount Sinai, New York, NY). Human colorectal carcinoma cells, HCT116 cells, were obtained from ATCC (#CCL-247, ATCC, Manassas, VA). Both cell types were cultured and expanded in tissue culture-treated plastic dishes. Cultures were passaged when cells reached 70-90% confluency. Both cell types were cultured with Dulbecco’s Minimum Essential Medium (DMEM, Millipore-Sigma, St. Louis, MO) containing 10% fetal bovine serum (FBS, Hyclone, Logan, UT). Cells were detached from the substrate with Trypsin/EDTA (Hyclone) and resuspended in media before use in organoid or spheroid fabrication.

### Organoid Fabrication

Spheroids of HCT116 cells (1.0×10^4^ cells each) were prepared by homogenously suspending cells in culture media at 1.0×10^5^ cells/mL followed by dispensing 100 µL of cell-media suspension into each well of an ultra-low attachment round-bottom 96-well plate (CoStar #7007, Corning, Corning, NY). Cells were observed each day for 3 days, by then spheroids had formed tight clusters without irregularity, and used immediately for organoid fabrication.

Organoids were fabricated as described recently (**Suppl. Fig.1a**).^9^ Type I Rat Tail Collagen (#354236, Corning) was prepared according to the manufacturer’s protocol at a concentration of 2mg/mL. LX2 cells were trypsinized and counted, then suspended in prepared collagen at 5.0×10^6^ cells/mL. Media from plates containing spheroids was aspirated, and 100 µL of LX2-collagen solution was pipetted into each spheroid well, carefully to avoid disturbing the spheroid structure. The LX2-collagen-spheroid mixture was slowly pipetted up and down to suspend the spheroid, then the whole volume was dispensed into a custom polydimethylsiloxane (DOW Sylgard 184, Midland, MI) mold ^9^ ensuring relatively central placement of the spheroid within the polymerizing hydrogel (30 min at 37ºC) (**Suppl. Fig.1a**). Upon complete collagen polymerization, media was slowly added and molds were removed. Organoids were cultured for varying durations depending on the scope of the experiment.

### Subcutaneous implantation

Six-week-old female athymic nude mice were obtained from Charles River Laboratories (Wilmington, MA). Animals were anesthetized using 2% isoflurane and given a pre-operative subcutaneous injection of 5 mg/kg ketoprofen. Two small skin incisions were made on the dorsal side approximately 15 mm from the midline. Subcutaneous pockets were generated using blunt dissection and implanted with one organoid per site. Incisions were closed with 4-0 vicryl suture (Ethicon, Somerville, NJ) and dressed with Tegaderm™ film (3M, Maplewood, MN). All procedures were performed in strict accordance with the NIH Guide for the Care and Use of Laboratory Animals, and the policies of the Wake Forest University Institutional Animal Care and Use Committee (IACUC) (Protocol No. A17-036). Animals were euthanized at 2 and 4 weeks by CO_2_ asphyxiation followed by cervical dislocation. Skin flaps were opened and organoid explants retrieved.

### Gross morphological assessment

Gross macroscopic images of *in vivo* organoid explants were taken with a smartphone camera (Samsung, Seoul, South Korea) with a metric ruler held within the frame of the image. ImageJ software (National Institutes of Health, Bethesda, MD) was used to assess explant size after initial calibration.

### Chemotherapy treatments

Organoids with embedded spheroids were cultured for 72 hours, then transferred to new well plates and incubated with media containing chemotherapeutics. Organoids were exposed to chemotherapeutics for a further 72 hours before analysis. Chemotherapeutic formulations were prepared using the following concentrations: 5-Fluorouracil 1mM, Oxaliplatin 25µM, Irinotecan 50µM, Leucovorin 50µM, and Regorafenib 50µM.

### Histological and immunohistochemical (IHC) analysis

Samples were fixed in 4% paraformaldehyde overnight at 4°C, then washed with phosphate buffered saline (PBS), and stored in 70% ethanol. Following paraffin processing and embedding, 5 µm serial sections were cut using a microtome (Leica Microsystems Inc., Buffalo Grove, IL) and mounted to slides. For all staining procedures, slides were baked for 1 h at 60ºC followed by standard deparaffinization and rehydration. Hematoxylin & Eosin (H&E) staining was performed by core facilities at the Wake Forest Institute for Regenerative Medicine. Picrosirius Red (PRS) staining was done using a commercial staining kit (#24901, PolySciences, Warrington, PA) following the manufacturer’s protocol. Masson’s Trichrome staining was performed using a commercial staining kit (#HT15, Millipore-Sigma) following the manufacturer’s protocol.

For immunohistochemistry (IHC), all incubations were performed at room temperature unless otherwise stated. Antigen retrieval was performed using Proteinase K (DAKO, Carpinteria, CA) incubation for 15 min. Samples were permeabilized with 0.05% Triton-X (insert vendor) in PBS for 5 min. Non-specific antigen blocking was performed using Protein Block Solution (#ab156024, Abcam, Cambridge, MA) incubation for 30 min. Slides were then incubated with the appropriate primary antibody against CK-18 (#ab82254, Abcam), FAK (#ab40794, Abcam), β-Catenin (#71-2700, Invitrogen-ThermoFisher), E-Cadherin (#ab40772, Abcam), N-Cadherin (#ab76011, Abcam), CD44 (#ab51037, Abcam), CD133 (#orb10288, biorbyt, San Francisco, CA) or Ki-67 (#ab16667, Abcam) at recommended dilutions in a humidified chamber overnight at 4°C. Next, slides were thoroughly washed and incubated for 1 h with the appropriate secondary antibody: biotinylated anti-rabbit (BA-1000, Vector Laboratories) or biotinylated anti-mouse (BA-2000, Vector Laboratories). Slides were then washed and incubated with VECTASTAIN ABC reagent (PK-4000, Vector Laboratories) for 30 min. Signal exposure was timed and visualized while slides were incubated with DAB (SK-4105, Vector Laboratories) or Vector Red (SK-5105, Vector Laboratories) substrate. Double-stained slides followed this protocol twice in succession for each individual marker. Vector Red substrate was used first and DAB second. Relevant control slides were prepared for each condition and each antibody combination by omitting the primary antibody incubation. Slides were mounted with MM24 (#3801120, Leica, Wetzlar, Germany), and light microscopy images, using linearly polarized light for PSR-stained sections, were captured with an Olympus BX63 microscope (Olympus, Center Valley, PA) with an Olympus DP80 camera (Olympus).

### Image analysis and quantification

Spheroid size was assessed using H&E-stained light micrographs and a MatLab (2018b, MathWorks, Natick, MA) script to quantify average diameter. Counts of infiltrative blood vessels and vessel size were quantified using a MatLab script. Images were captured with an Olympus IX83 microscope (Olympus, Center Valley, PA) equipped with a Hamamatsu Orca-Flash 4.0 camera (Hamamatsu, Shizuoka, Japan).

PRS signal was quantified in two ways: hue analysis of collagen signal and collagen fiber geometric parameter segmentation. Hue analysis to identify different levels of collagen bundling and fibrilization was performed using a MatLab script. Fiber parameter segmentation and quantification was performed using CT-FIRE (Laboratory for Optical and Computation Instrumentation, University of Wisconsin).^21^

Masson’s Trichrome signal hue was quantified similarly to PRS hue analysis. Red and blue pixels were identified and counted using a MatLab script. Ratios of colored pixels versus total tissue pixels were then calculated for comparison.

IHC images were imported as uncompressed files into Visiopharm software (Broomfield, CO) for analysis and quantification. In samples where tumor cells were difficult to distinguish from stromal cells (*in vitro* and *in vivo* implanted organoids), a script was written to deconvolve Vector Red signal, then isolate Vector Red stained CK-18 positive cells which specifically labels HCT116 cells. After HCT116 cells were segmented, a second script was written to deconvolve DAB signal and quantify the expression or localization of: β-catenin, E-Cadherin, FAK, Ki67, and N-Cadherin. For β-catenin and Ki67, nuclei were marked as positive or negative for DAB staining and counted. For E-Cadherin and N-Cadherin, cells were marked positive if they had complete membrane localization of DAB and negative if not. For FAK, ratios of total tumor area that stained positive for DAB were generated.

### Statistical Analysis

All experiments were performed using n=3 unless otherwise stated. Statistical analysis of parametric data was performed using Student’s t-test or one-way analysis of variance (ANOVA) with Tukey’s multiple comparison *post hoc* test. Statistical analysis of non-parametric data was performed using Kolmogorov-Smirnov chi-square tests. Significance was defined as α ≤ 0.05 and all p-values are reported with their respective data-sets. GraphPad Prism software v6.0 (GraphPad Software, La Jolla, CA) was used for all analyses.

## Author contributions

M.D., S.H., A.S., and S.S. designed experiments. M.D. performed all *in vitro* experiments. M.D., S.H., and A.D. performed all *in vivo* experiments. M.D., A.D., and E.W. performed immunohistochemical staining and imaging. M.D. designed and performed all digital segmentation analysis. M.D. designed all figures. M.D., S.H., and S.S. prepared manuscript text. All authors reviewed data and text.

## Acknowledgments

We thank Dr. Scott Friedman from the Icahn School of Medicine at Mount Sinai, New York, for providing LX2 cells.

