## Supplemental Information for "Simulating the human tumor microenvironment in colorectal cancer organoids *in vitro* and *in vivo*"

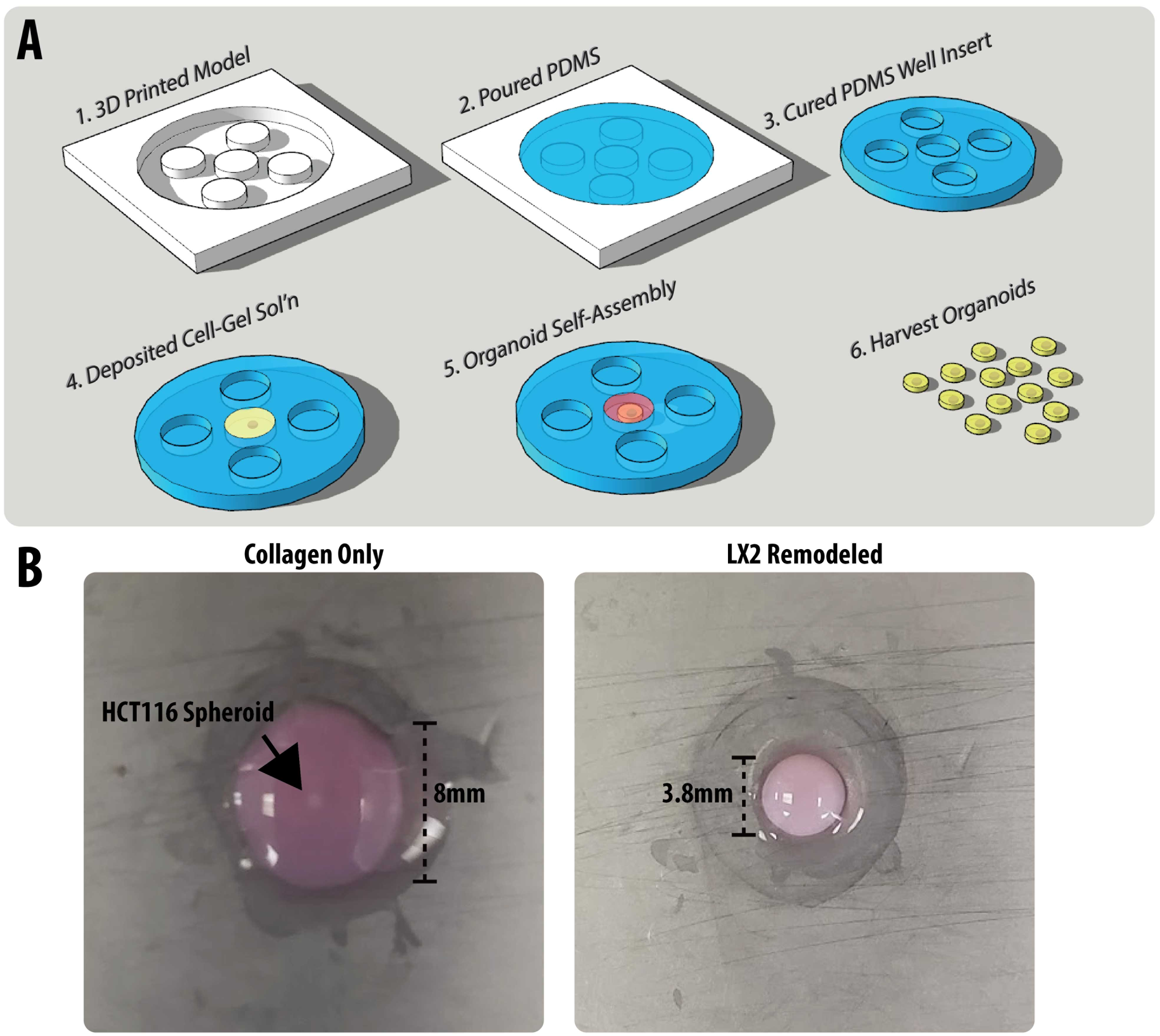

**Supplementary Figure 1: Organoid fabrication and gross morphology.** (**A**) Organoid molds were produced through additive printing of a negative mold, then deposited silicone was cured to yield a 6-well plate microwell insert. Then cell-hydrogel solution was deposited, with a suspended HCT-116 spheroid, into microwells and allowed to gelate, then self-assemble over experimental timeline before being harvested for further analysis. (**B**) LX2 organoids undergo size contraction compared to collagen-only controls.

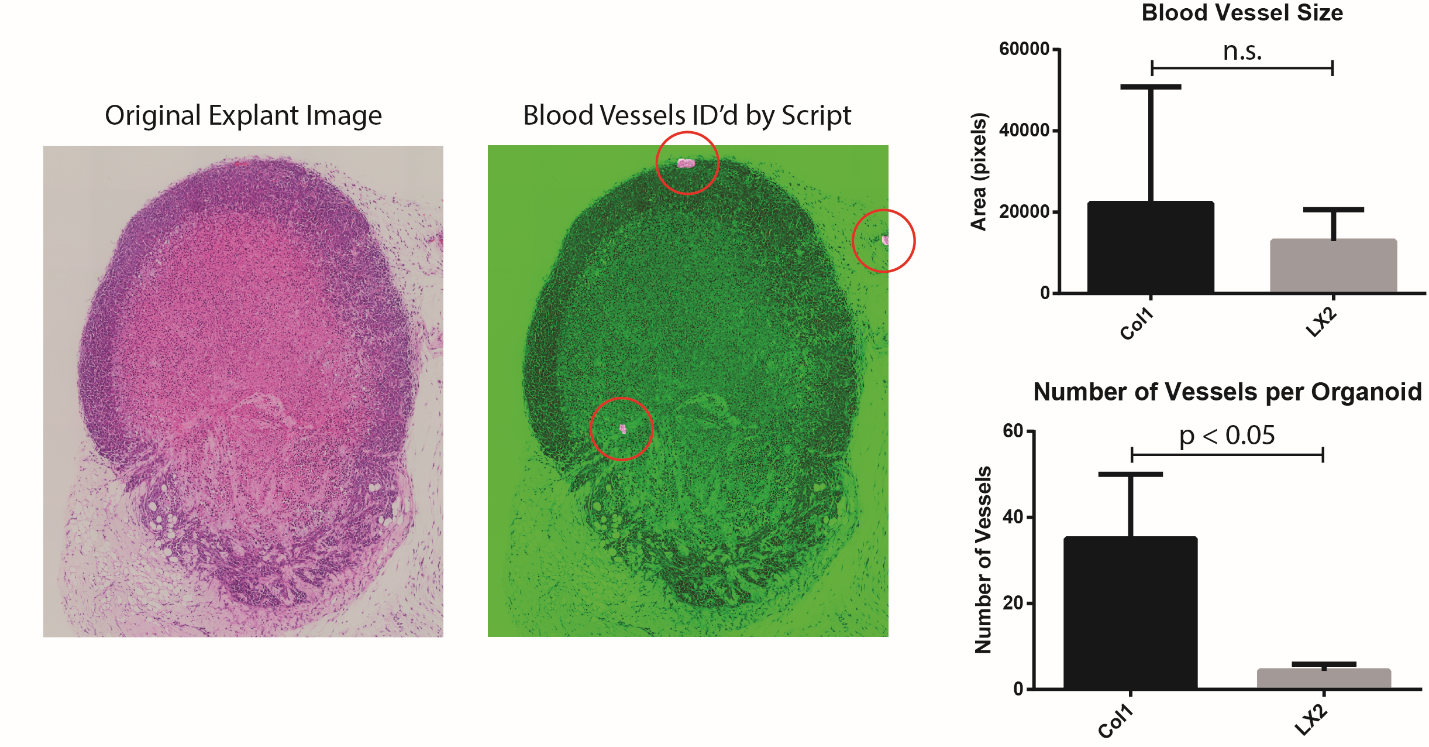

**Supplementary Figure 2: Blood vessel quantification.** H&E-stained micrographs were used for analysis of blood vessel quantity and size. A MatLab script identifies vessel structures, then size and number of vessels were quantified.

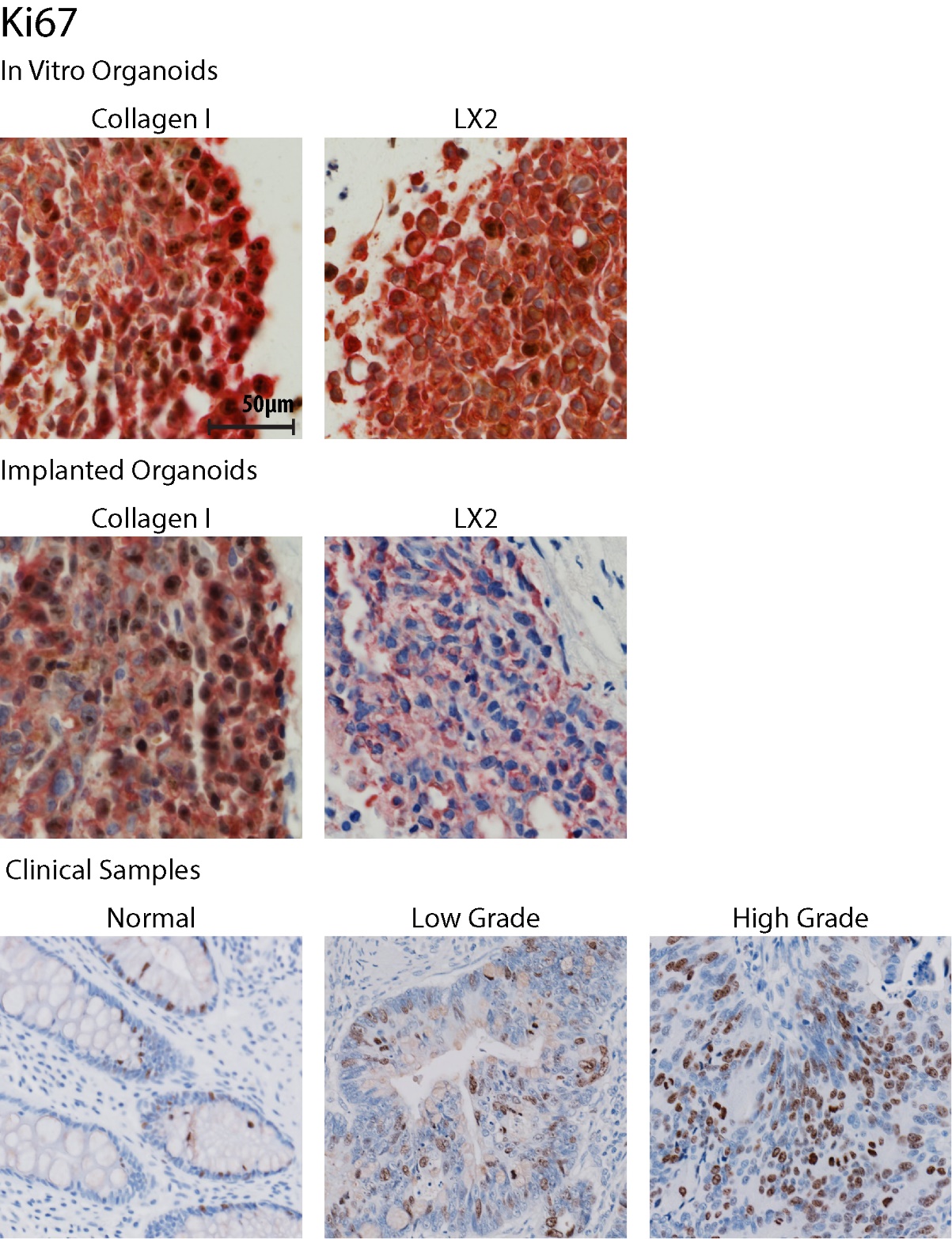

**Supplementary Figure 3: Ki67 immunohistochemical staining.** Both *in vitro* and *in vivo* organoids were double stained (DAB and Vector Red) to identify HCT-116 cells from LX2 and mouse cells.

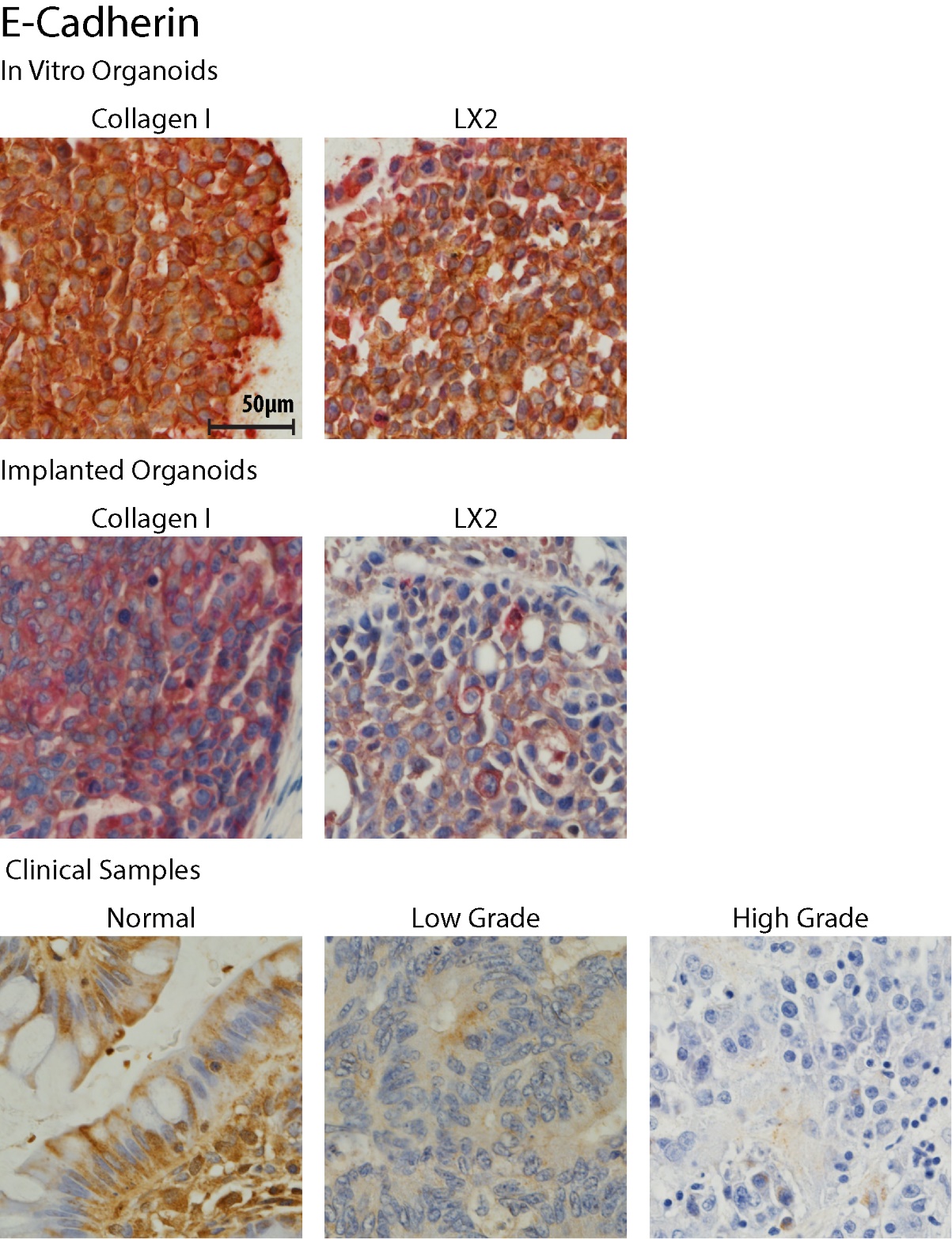

**Supplementary Figure 4: E-Cadherin immunohistochemical staining.** Both *in vitro* and *in vivo* organoids were double stained (DAB and Vector Red) to identify HCT-116 cells from LX2 and mouse cells.

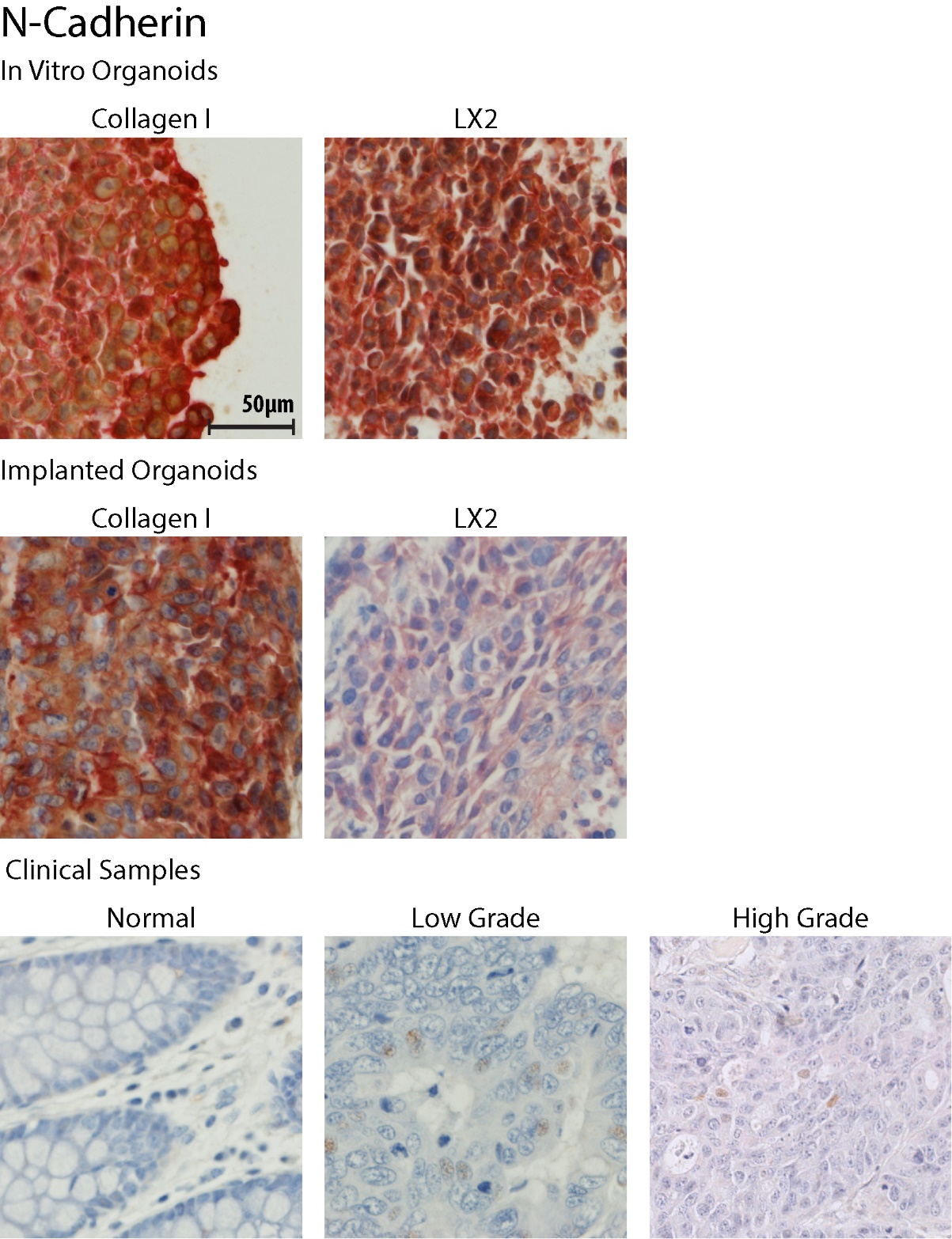

**Supplementary Figure 5: N-Cadherin immunohistochemical staining.** Both *in vitro* and *in vivo* organoids were double stained (DAB and Vector Red) to identify HCT-116 cells from LX2 and mouse cells.

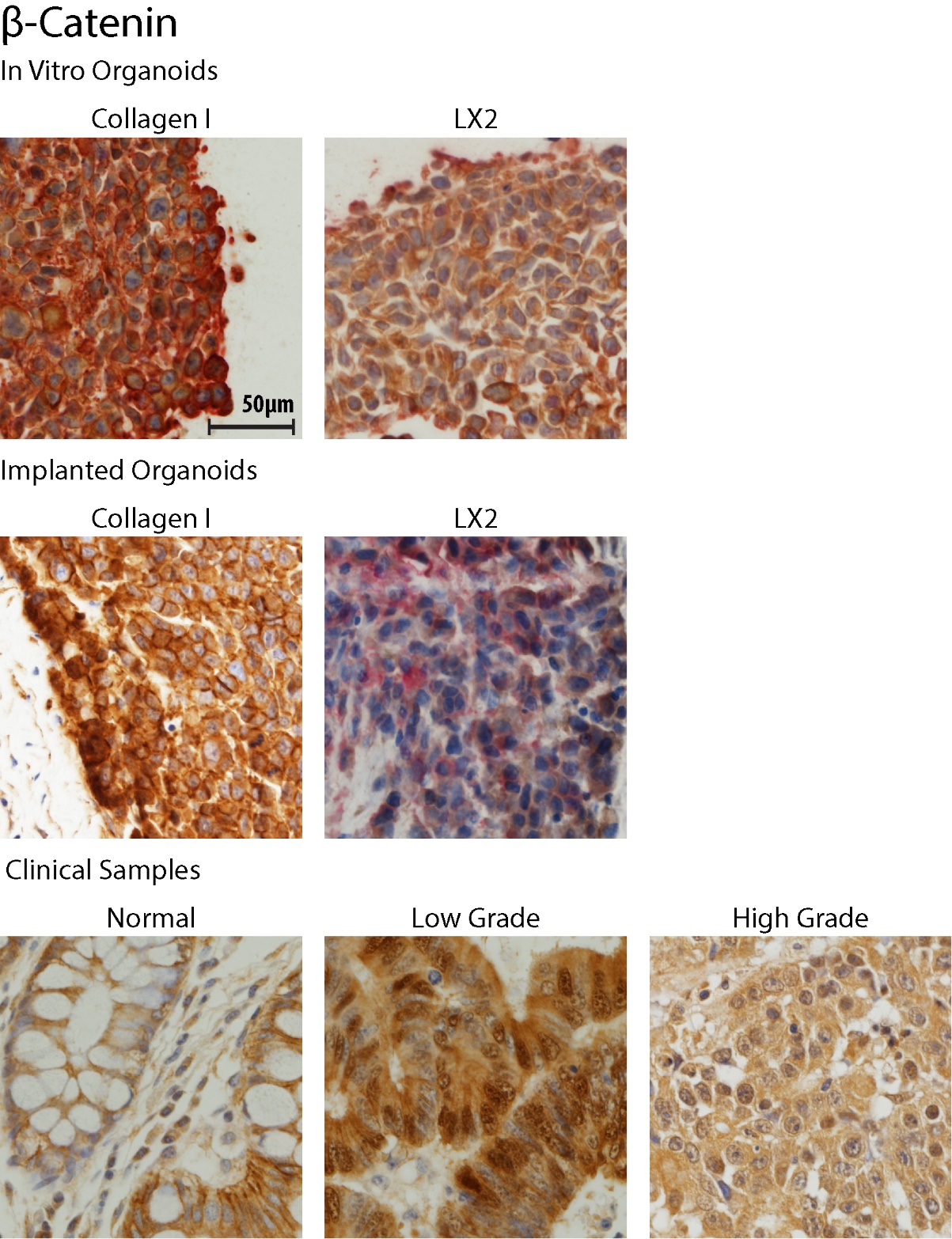

**Supplementary Figure 6: β-Catenin immunohistochemical staining.** Both *in vitro* and *in vivo* organoids were double stained (DAB and Vector Red) to identify HCT-116 cells from LX2 and mouse cells.

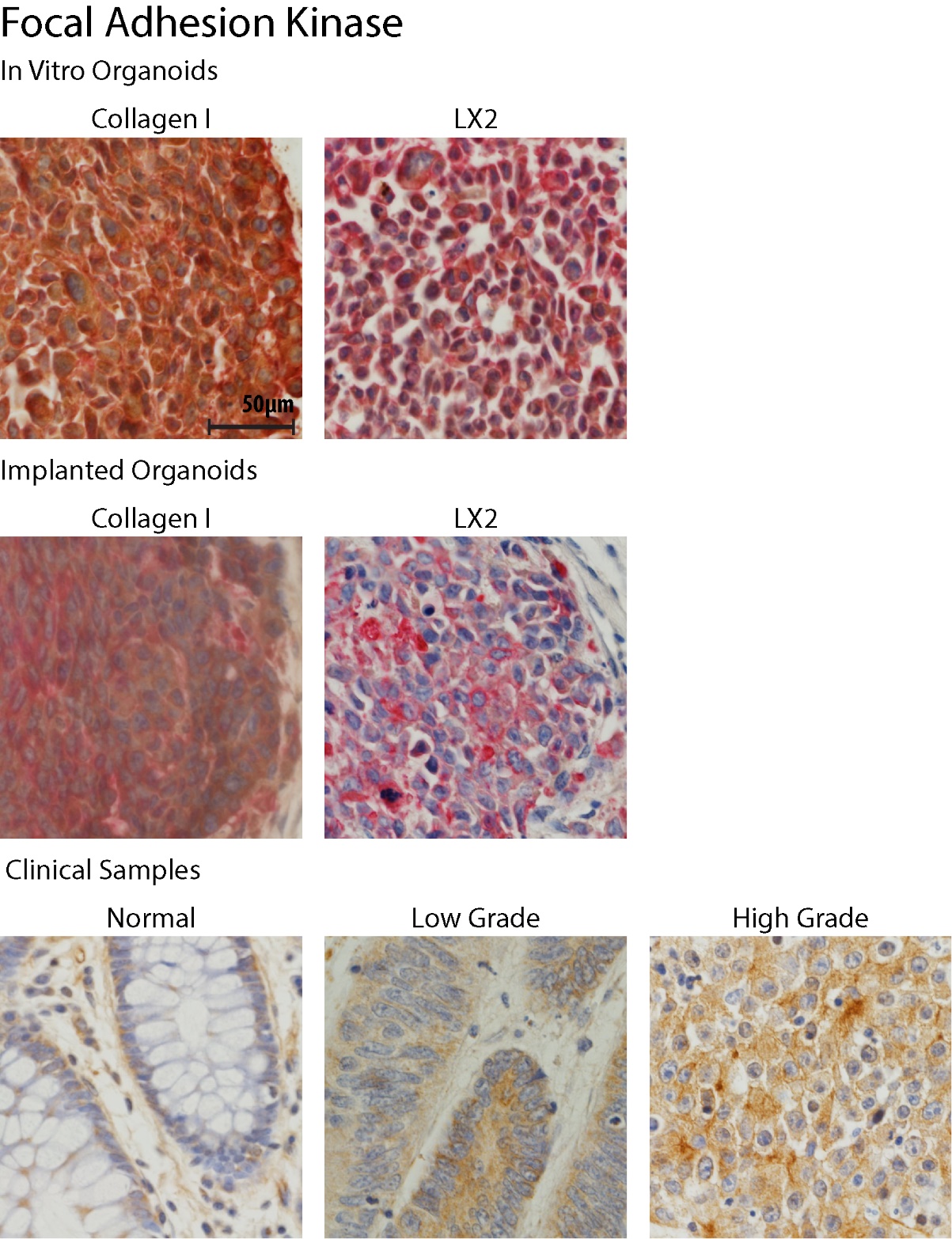

**Supplementary Figure 7: FAK immunohistochemical staining.** Both *in vitro* and *in vivo* organoids were double stained (DAB and Vector Red) to identify HCT-116 cells from LX2 and mouse cells.

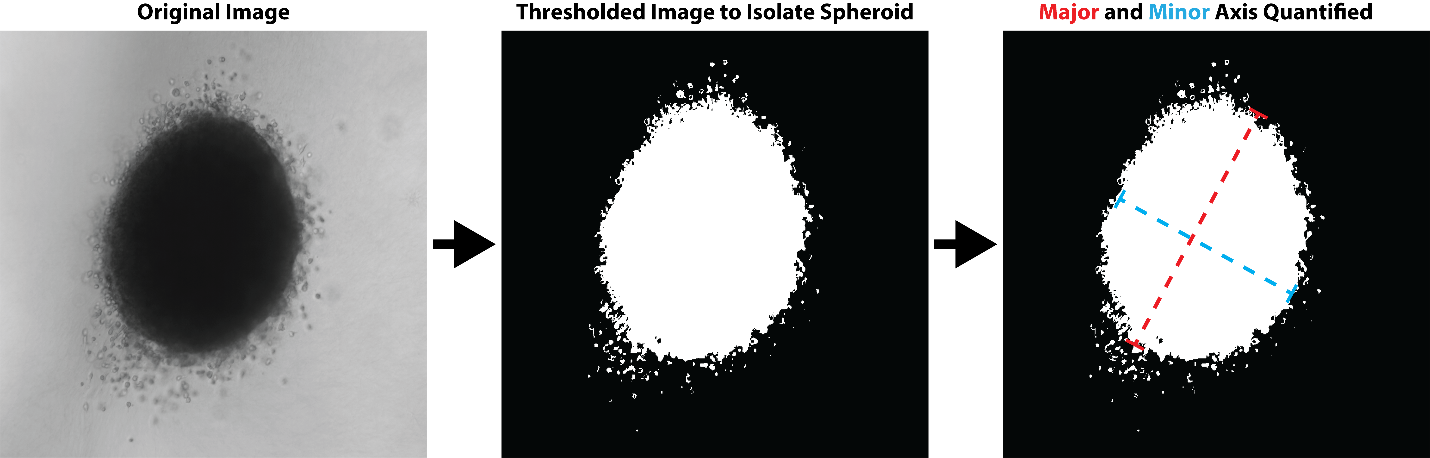

**Supplementary Figure 8: Quantification of spheroid size after organoid culture.** Organoids were imaged under brightfield conditions to visualize the spheroid body. A MatLab script was then used to segment the spheroid and quantify average diameter.

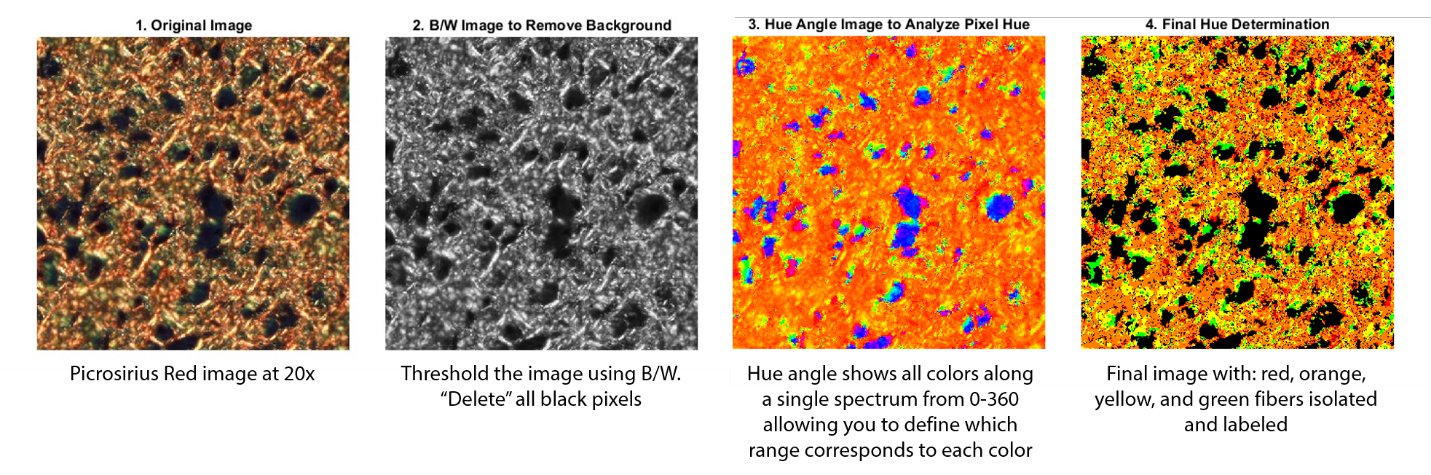

**Supplementary Figure 9: Picrosirius red hue analysis.** PSR-stained images were captured under polarized light, then a MatLab script was used to quantify pixels of varying hue signal.

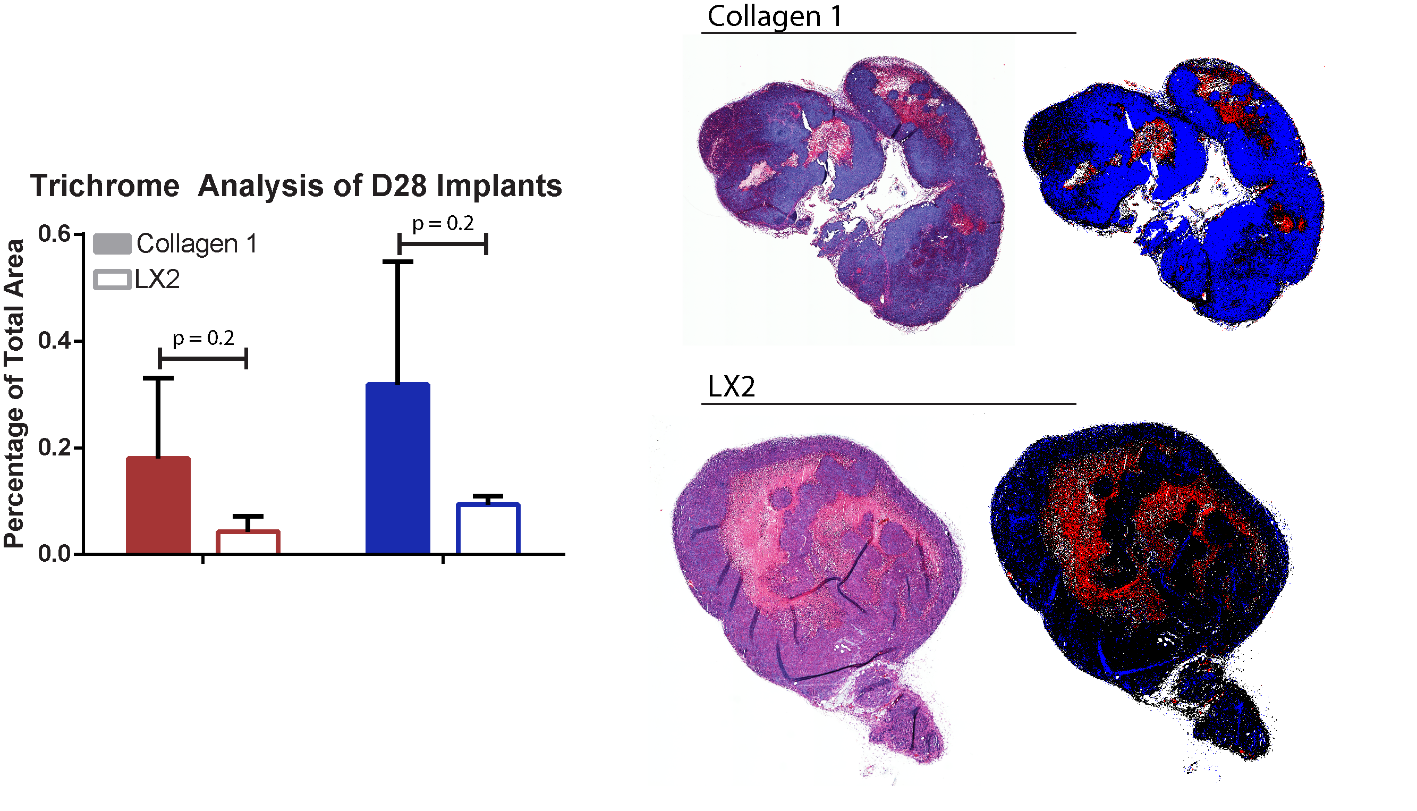

**Supplementary Figure 10: Trichrome staining and analysis.** *In vivo* implants were stained with Masson’s Trichrome and bright field images were analyzed using a MatLab script to segment blue and red pixels.
